# Choosing the optimal population for a genome-wide association study: a simulation using whole-genome sequences from rice

**DOI:** 10.1101/798850

**Authors:** Kosuke Hamazaki, Hiromi Kajiya-Kanegae, Masanori Yamasaki, Kaworu Ebana, Shiori Yabe, Hiroshi Nakagawa, Hiroyoshi Iwata

## Abstract

A genome-wide association study (GWAS) needs to have a suitable population. The factors that affect a GWAS, e.g. population structure, sample size, and sequence analysis and field testing costs, need to be considered. Mixture populations containing subpopulations of different genetic backgrounds may be suitable populations. We conducted simulation experiments to see if a population with high genetic diversity, e.g., a diversity panel, should be added to a target population, especially when the target population harbors small genetic diversity. The target population was 112 accessions of *Oryza sativa* subsp. *japonica*, mainly developed in Japan. We combined the target population with three populations that had higher genetic diversities. These were 100 *indica* accessions, 100 *japonica* accessions, and 100 accessions with various genetic backgrounds. The results showed that the GWAS power with a mixture population was generally higher than with a separate population. Also, the GWAS optimal population varied depending on the fixation index *F*_*ST*_ of the quantitative trait nucleotide (QTN) and its polymorphism of QTN in each population. When a QTN is polymorphic in a target population, a target population combined with a higher diversity population improves the QTN detection power. Investigating *F*_*ST*_ and the expected heterozygosity *H*_*e*_ as factors influencing the detection power, we showed that SNPs with high *F*_*ST*_ or low *H*_*e*_ are less likely to be detected by GWAS with mixture populations. Sequenced/genotyped germplasm collections can improve the GWAS detection power by using a subset of them with a target population.

Core ideas (3-5 impact statements, 85 char max for each)

- Genome-wide association studies with mixture populations are expected to improve the detection power of novel genes due to the increase of the sample size although the influence of population structure is a concern.
- When a quantitative trait nucleotide (QTN) is polymorphic in a target population, a combination of the target population and a population with higher diversity than the target population improves the detection power of the QTN.
- We found that the fixation index (*F*_*ST*_) and the expected heterozygosity (*H*_*e*_) were strongly related to the detection power of QTNs.
- Germplasm collections which have been already sequenced/genotyped are useful for improving the detection power of GWAS without any addition of sequence costs by using a subset of them with a target population.

## INTRODUCTION

Recently, as genome sequencing costs have continued to decrease (Metzker, 2010), the whole-genome sequences of a large number of cultivars/lines have become available for major crop species, such as rice (Li et al., 2014; Wang et al., 2018). A genome-wide association study (GWAS) based on whole-genome sequences can more efficiently and accurately identify genes that control important agronomic traits than previous methods (Koboldt et al., 2013; Ott et al., 2015; Yano et al., 2016).

It is important to prepare an appropriate population to be analyzed when attempting to detect candidate genes using GWAS techniques. For example, to avoid potential false positives caused by population stratification/structure, a GWAS population should be selected that results in low stratification (Begum et al., 2015; Yano et al., 2016). However, if such a population is selected as an analytical population for a GWAS, the sample size may be limited and the detection power of the GWAS will decrease (Korte and Farlow, 2013). Therefore, when designing an appropriate GWAS population, one should be aware of the trade-off relationship between population stratification and sample size.

When preparing the population to be analyzed, the factors that directly affect the GWAS results, such as population structure, sample size, and the sequence analysis and cultivation testing costs, need to be considered. In recent years, the whole-genome sequences of a large number of cultivars/lines have become publicly available due to highly efficient sequencing analyses and database enrichment. The publically available whole-genome sequence data will improve GWASs and could enable researchers to avoid the costs of sequencing analyses. For example, in rice, “The 3,000 Rice Genomes Project” (Li et al., 2014; Wang et al., 2018) by the International Rice Research Institute (IRRI) is a well-known whole-genome sequence dataset that is available in the “Rice SNP-Seek Database” (Alexandrov et al., 2015; Mansueto et al., 2016; 2017). Therefore, an appropriate GWAS population could potentially utilize existing public sequence data.

A mixture population obtained by mixing subpopulations with different genetic backgrounds could also potentially be used in a GWAS. An advantage of using such a mixture population is that it should improve the detection of causal variants by increasing the sample size. Conversely, a GWAS with a mixture population may suffer from large numbers of false positives caused by the population structure. Although a mixed effect model that suppresses the influence of the population structure has been proposed (Yu et al., 2006), such a mixture population has rarely been analyzed by a GWAS.

An actual data analysis of rice using whole-genome sequences showed that the detection power of a GWAS improved when *Oryza sativa* subsp. *japonica* and *Oryza sativa* subsp. *indica* populations were combined (Misra et al., 2017). Furthermore, the identification of new rice genes using a GWAS and populations with extremely high genetic diversities has also been previously reported (Zhao et al., 2011). Conversely, it has been reported that the genetic differentiation between subpopulations in a population with high genetic diversity could cause a reduction in the power of a GWAS (Huang et al., 2012). Therefore, real data studies have been inconsistent about whether mixture populations or populations with high genetic diversities should be used in a GWAS. However, these previous studies mostly analyzed actual data, and there have been no theoretical simulation studies that have considered the possibility of using a mixture population in a GWAS. Furthermore, no previous studies have discussed which kinds of populations should be mixed to improve the GWAS detection power or which kinds of populations are most appropriate for a GWAS. Therefore, in this study, we conducted simulation experiments to see whether adding a population with a high genetic diversity compared to a target population (e.g., adding a diversity panel to a target population) is appropriate, especially when the genetic diversity of the target population is small.

## MATERIALS AND METHODS

### Materials (populations used in the GWAS)

In this study, 112 accessions of *Oryza sativa* subsp. *japonica* (referred to as “A”), which were accessions that had mainly been developed in Japan, were used as a target population with low genetic diversity (Yabe et al., 2016). We used the following three populations selected from “The 3,000 Rice Genomes Project” (J. Y. Li et al., 2014), i.e., 100 accessions of *Oryza sativa* subsp. *indica* (referred to as “B”), 100 accessions of *Oryza sativa* subsp. *japonica* (referred to as “C” or temperate), and 100 accessions of *Oryza sativa* with various genetic backgrounds (referred to as “D” or diverse), as populations with higher diversities than the target population (Table S1 in Supplemental File 1). The process used to select each of the 100 accessions is described in Supplemental File 9. One accession (IRIS ID: IRIS 313-11868) was duplicated in populations B and D. Among populations B, C, and D, the B population was the most differentiated from A, whereas C was the most similar to A. Population D contained subsp. *indica*, subsp. *japonica*, and *aus*, and *aromatic* rice accessions, which meant that the D population had the highest genetic diversity. Fig. 1 is an unrooted phylogenetic tree that shows the genetic relationships among accessions belonging to populations A, B, C, and D.

**Fig. 1.**
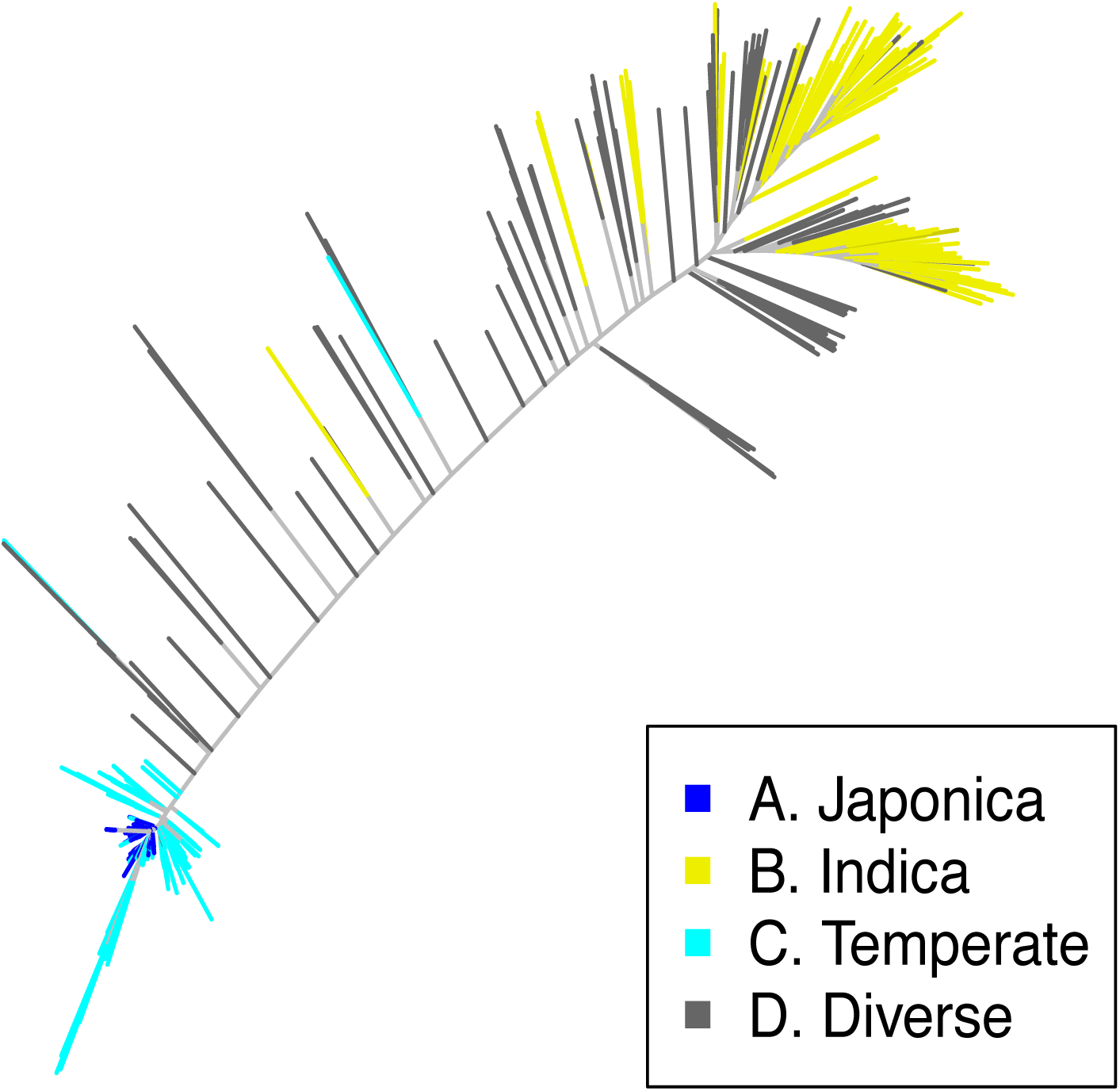
Unrooted phylogenetic tree plot for four non-mixture populations. Unrooted phylogenetic tree plot for the four non-mixture populations, which consisted of 112 accessions of *japonica* (A), 100 accessions of *indica* (B), 100 accessions of temperate *japonica* (C), and 100 diverse accessions (D) with neighbor-joining method.

The genetic relationships among the accessions were estimated by the neighbor-joining (NJ) method (Saitou and Nei, 1987) using the R package “ape” version 5.3 (Paradis et al., 2004). The genetic distances were estimated according to the Jukes and Cantor (1969) model. In addition to these four populations, we synthesized three populations by combining population A with populations B, C, or D. The mixture populations A + B, A + C, and A + D were named “E”, “F”, and “G”, respectively. We compared the QTN detection power the GWAS when the seven non-mixture (A, B, C, and D) and mixture populations (E, F, and G) were used.

### Genotype data

Whole genome sequencing data were available for the accessions (Jarquin et al., 2019). Details about the DNA extraction and whole genome sequencing techniques are provided in a previous report (Jarquin et al., 2019). The data sets deposited in the DDBJ Sequence Read Archive (SRA106223, ERA358140, DRA000158, DRA000307, DRA000897, DRA000927, DRA007273, DRA007256, and DRA008071) were reanalyzed. We processed the whole-genome sequence data as follows so that they could be used in the GWAS. Adapters and low-quality bases were removed from paired reads using the Trimmomatic v0.36 program (Bolger et al., 2014). The preprocessed reads were aligned using Os-Nipponbare-Reference-IRGSP-1.0 (Kawahara et al., 2013) and the bwa-0.7.12 mem algorithm with the default options (H. Li, 2012). The binary alignment map (BAM) files deposited in the Rice SNP-Seek database were also reanalyzed. Single nucleotide polymorphism (SNP) calling was based on alignments determined using the Genome Analysis Toolkit (GATK), 3.7-0-gcfedb67 (McKenna et al., 2009; Auwera et al., 2014) and Picard package V2.5.0 (http://broadinstitute.github.io/picard). The mapped reads were realigned using RealignerTargetCreator and indelRealigner in the GATK software. The SNPs and InDels were called at the population level using the UnifiedGenotyper in GATK and the -glm BOTH option. We extracted bi-allelic sites in all the accessions from the variants using VCFtools version 0.1.13 (Danecek et al., 2011). Then, imputations were imputed using Beagle version 4.1 (Browning and Browning, 2016). Finally, we analyzed the SNPs with minor allele frequencies (MAFs) ≥ 0.05 in each population. In the analysis, the genotypes were represented as -1 (homozygous of the reference allele), 1 (homozygous of the alternative allele) or 0 (heterozygous of the reference and alternative alleles). Out of all the whole-genome sequence polymorphisms, only the SNPs on chromosome 1 were analyzed. The number of SNPs on chromosome 1 in each population is shown in Table 1.

**Table 1.**
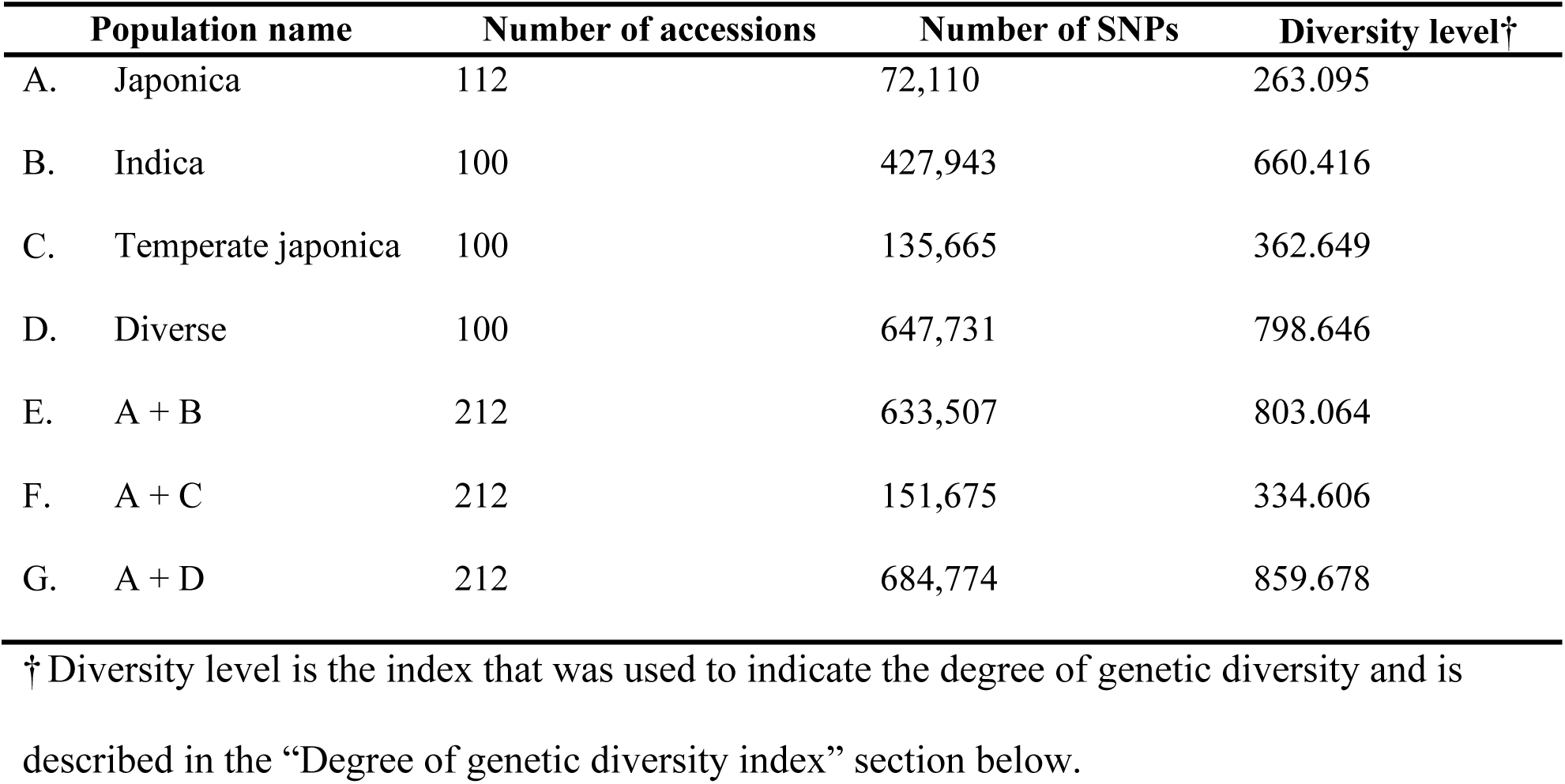
Number of SNPs and the diversity level of non-mixture and mixture populations.

### Generating phenotype data

Phenotypic data were simulated using the following formula:

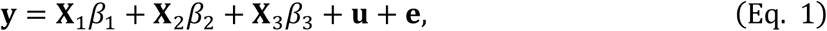

where **y** is the vector that represents the simulated phenotypic values for all 411 accessions; **X** is the design matrix representing the genotypes of three quantitative trait nucleotides (QTNs) with scores -1, 0, or 1; **β** = [*β*_1_ *β*_2_ *β*_3_]^T^ is the vector representing the effects of the three QTNs, **u** is the vector for polygenetic effects, and **e** is the residuals vector. Three QTN-SNPs whose MAF was equal to or larger than 0.05 in all 411 accessions (672,923 SNPs in total) were randomly selected from the SNPs on chromosome 1. The simulations were divided into five categories (low, lower-middle, middle, higher-middle, high) based on the fixation index (*F*_*ST*_) between populations A and B for the first QTN (Fig. S1 in Supplemental File 2). We assumed that the first QTN had four times greater variance than the remaining two QTNs (referred to as “QTN1”, “QTN2”, and “QTN3” respectively). The remaining two QTNs were chosen randomly from SNPs where the *F*_*ST*_ between A and B were low (SNPs whose *F*_*ST*_ value was in the lower 20% category among the 672,923 SNPs). The *F*_*ST*_ for each marker was calculated according to Wright (1965) as follows:

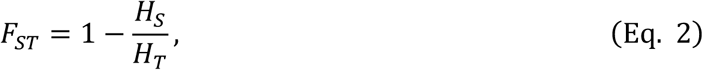

where *H*_*S*_ is the average of the expected heterozygosity based on the allele frequencies of populations A and B, and *H*_*T*_ is the expected heterozygosity based on the average allele frequency of populations A and B. *H*_*S*_ and *H*_*T*_ were calculated as follows:

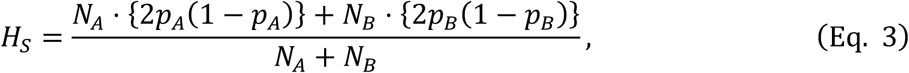

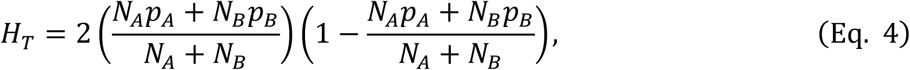

where *p*_*A*_, *p*_*B*_, *N*_*A*_, and *N*_*B*_ are the allele frequencies and the sample sizes of populations A and B respectively, and *N*_*A*_ = 112 and *N*_*B*_ = 100. The *F*_*ST*_ distribution between A and B is shown in Fig. S1, which also shows the thresholds for the five *F*_*ST*_ categories.

The polygenetic effect in Eq. 5 was sampled from the multivariate normal distribution whose variance-covariance matrix was proportional to the additive numerator relationship matrix **A** and was normalized so that their variance was equal to that of the three QTN effects.

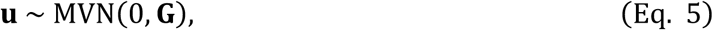

where 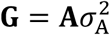 is the genetic covariance matrix, and the additive genetic variance 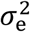 was automatically determined from the relationship with heritability. In this study, the additive numerator relationship matrix **A** was estimated based on the marker genotype data for 402,509 SNPs, which consisted of the core SNPs (defined by the Rice SNP-Seek Database as the “404k CoreSNP Dataset”) in all 12 chromosomes (this marker genotype data was prepared separately from the whole-genome sequence data), using the “A.mat” function in R package “rrBLUP” version 4.5 (Endelman and Jannink, 2012; Endelman, 2011).

The residual **e** in Eq. 6 was sampled identically and independently from the normal distribution, and was then normalized so that the narrow-sense heritability was equal to 0.6. Residual **e** was calculated using the following formula:

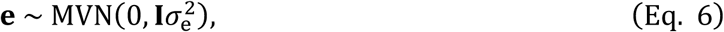

where **I** is an identity matrix, and the residual variance 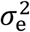 was determined so that the heritability was equal to 0.6.

### Genome-wide association study (GWAS) using simulated data

We performed a GWAS on the seven non-mixture (A, B, C, D) and mixture populations (E, F, and G) using the marker genotype data and the simulated phenotypic data. We fitted the linear mixed model (Yu et al., 2006).

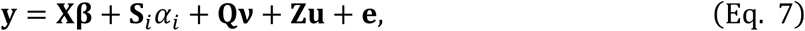

where **y** is the vector of phenotypic values, **Xβ, S**_*i*_*α*_*i*_, and **Q*v*** are the fixed effects terms, **Zu** is the random effects term, and **e** is the residuals vector. **β** represents all of the fixed effects other than **S**_*i*_***α***_*i*_, and **Q*v***, and X is the incidence design matrix corresponding to **β**. In this study, **Xβ** was an intercept. **S**_*i*_*α*_*i*_ is composed of **S**_*i*_, which is the *i*th marker of the genotype data, and *α*_*i*_, which is the effect of that marker. **Q*v*** is the term used to correct for the effect of population structure, and in this study **Q** was the matrix of two eigenvectors corresponding to the upper two eigenvalues of the additive numerator for relationship matrix **A**, Finally, **u** represents the polygenetic effects, and **Z** is the incidence design matrix corresponding to **u**.

We used the EMMAX and P3D algorithms to reduce the computation time (Kennedy et al., 1992; Kang et al., 2008; 2010; Zhang et al., 2010). The “GWAS” function in R package “rrBLUP” version 4.5 (Endelman, 2011) was used to perform the GWAS described above.

### Evaluation of the simulation results

The *p*-value (or −log_10_(*p*)) for each marker effect was estimated 100 times by the GWAS in five patterns according to the size of the *F*_*ST*_ for the seven non-mixture/mixture populations. In this study, the following summary statistics were mainly used to evaluate the GWAS results.

In the 100 simulations, the QTNs were not always polymorphic in each population (because the MAF of the whole population did not necessarily match the MAF of each individual population). In such cases, the −log_10_(*p*) value of a QTN that was not polymorphic within a population could not be calculated. Therefore, when two SNPs were polymorphic within that population and were adjacent to the QTN, then the statistic of the more significant SNP was used as the QTN statistic. Since it was difficult to detect such QTNs using a GWAS, we calculated the summary statistics by dividing two patterns depending on polymorphism patterns of QTN1, i.e., whether using all simulation results or using only results whose QTN1 was polymorphic in the target population (referred to as “All” and “Polymorphic in the population”, respectively).

### Correct detection rate (CDR) and −log_10_(*p*)

The first summary statistic was whether the −log_10_(*p*) rate for each QTN exceeded the threshold in each GWAS (referred to as “CDR; correct detection rate”). We assumed that QTNs would be successfully detected by the GWAS when the CDR was large. The −log_10_(*p*) value whose false discovery rate (FDR) was 0.05 was set as the threshold using the Benjamini-Hochberg method (Benjamini and Hochberg, 1995; Storey and Tibshirani, 2003). As the second summary statistic, we used the −log_10_(*p*) for each QTN in each GWAS, and we also assumed that QTNs were successfully detected by the GWAS when this statistic was large.

### Area under the curve (AUC)

We also regarded the mean of the AUC as one summary statistic. The AUC refers to the area under the receiver operating characteristic (ROC) curve (Fig. S2 in Supplemental File 3), which was obtained by plotting the false positive rate on the horizontal axis and the true positive rate on the vertical axis when the threshold was varied (Hanley and McNeil, 1982). The AUC was calculated using the following formula:

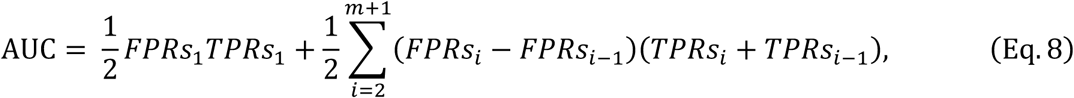

where *m* is the number of QTNs, and *m* = 3 in this study. The *FPRs* and *TPRs* are the *m* + 1 vectors whose *i*th elements are *FPRs*_*i*_ and *TPRs*_*i*_, respectively. *FPRs*_*i*_ = *TPRs*_*i*_ = 1 when *i* = *m* + 1. When 1 ≦ *i* ≦ *m*, the *FPRs*_*i*_ and *TPRs*_*i*_ represent the false positive rate and the true positive rate at the time when *i* QTNs exceed the threshold, respectively. They were calculated using the following formula:

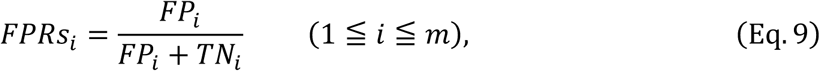

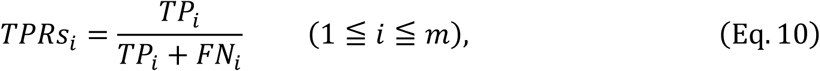

where *TP*_*i*_, *FP*_*i*_, *FN*_*i*_, and *TN*_*i*_ are the numbers of SNPs that are the true positives (where the SNP is a QTN and exceeds the threshold), the false positives (where the SNP is not a QTN but exceeds the threshold), the false negatives (where the SNP is a QTN but does not exceed the threshold), and the true negatives (where the SNP is not a QTN and does not exceed the threshold) at the time when *i* QTNs exceed the threshold respectively. When we evaluated the true/false positive rate, we considered the existence of linkage disequilibrium (LD) by investigating SNPs with LD as one set. In this study, we defined SNPs that satisfied the conditions that they were within 300 kb from the focused SNP and the condition that their squares of the correlation coefficients with the focused SNP were 0.35 or more as one set when considering LD. When we counted *TP*_*i*_, *FP*_*i*_, *FN*_*i*_, and *TN*_*i*_, we counted the number of the sets described above instead of the number of SNPs. The value for AUC calculated in this manner takes a value between 0 and 1. The GWAS is more successful when the AUC is closer to 1. Using the mean of the AUC as one of the summary statistics meant that it was possible to focus on each QTN and evaluate the overall results of the GWAS.

### Precision, recall, and F-measure

We calculated the mean of precision, the mean of recall, and the mean of the *F*-measure as other summary statistics to evaluate the GWAS results. These summary statistics can be calculated from the numbers of true positives, false positives, false negatives, and true negatives. More specifically, the precision can be calculated using the following formula:

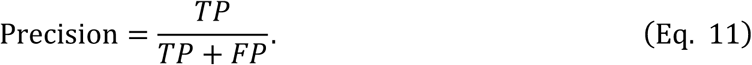

We regarded an SNP as “positive” when the −log_10_(*p*) of that SNP exceeded the threshold described above. The precision represents the ratio of the detected SNPs that were QTNs. The recall was defined using the following formula:

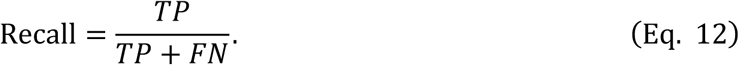

The recall represents the proportion of QTNs detected by the GWAS. Finally, the *F*-measure was calculated as the harmonic mean of the precision and the recall, and can be used to comprehensively evaluate the GWAS results. The *F*-measure was calculated using the following formula:

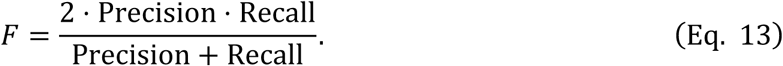

The greater these summary statistics, the more accurate the GWAS results were.

### Degree of genetic diversity index

In order to evaluate the relationship between genetic diversity and the CDR results, we prepared an index that indicated the degree of genetic diversity in each population. The Euclidean distance matrix between accessions for each population was calculated. The median for the off-diagonal elements of the distance matrix was used to indicate the degree of genetic diversity (referred to as the “diversity level”, Table 1). The median was chosen as the diversity level because the distribution of the distances between the accessions for E and G had a double peak. This was because, for mixture populations such as E and G, the distance within the subpopulations was small whereas the distance between subpopulations was large. Therefore, if the mean of the distances (almost the same as Nei’s gene diversity index (NEI, 1973)) is chosen as the diversity level, then there is a risk of overestimating the diversity level.

## RESULTS

### Comparisons between the CDR and AUC for the QTN1s in each population

The CDRs of the QTN1s in each population were calculated under ten conditions: five levels of *F*_*ST*_ between A and B and two patterns of QTN polymorphism, i.e., whether the QTN was polymorphic or not in the target population (Fig. 2 and Table S2 in Supplemental File 5).

**Fig. 2.**
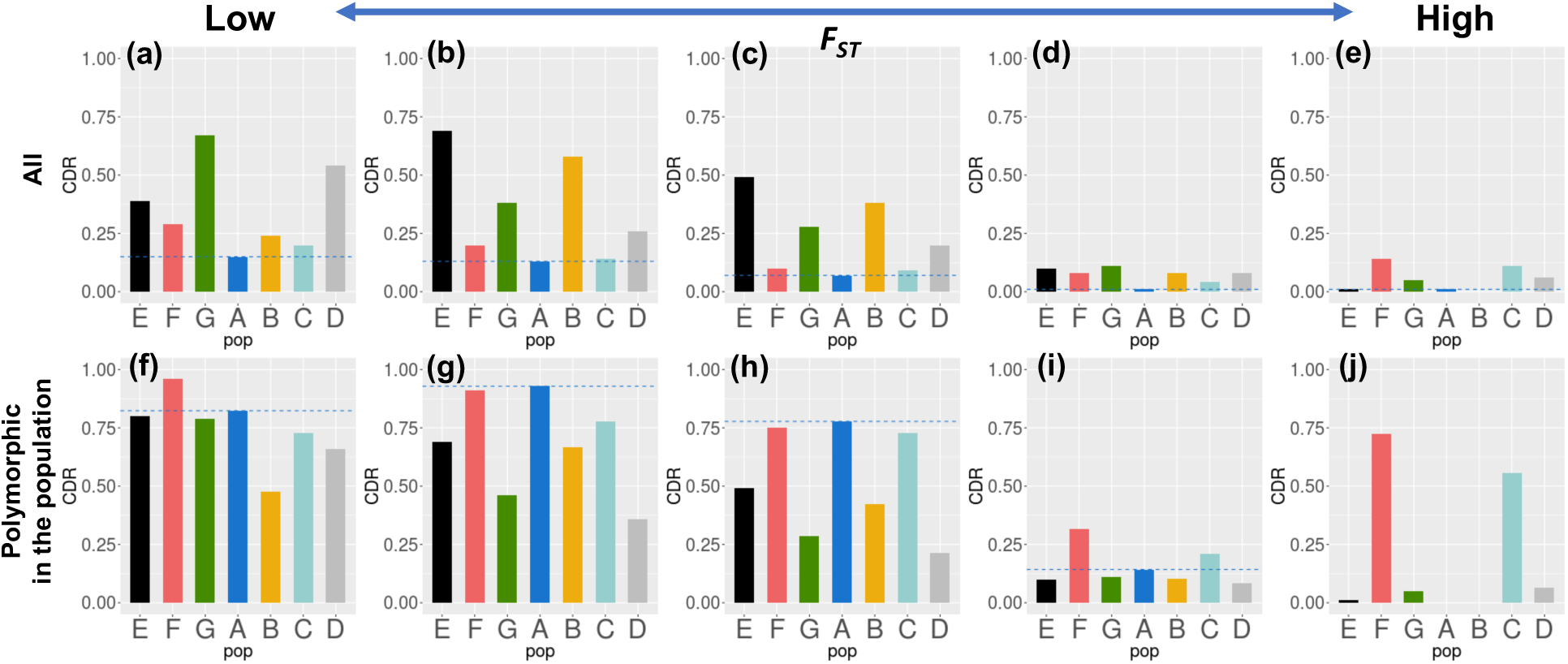
Correct detection rate for QTN1 in each population under ten conditions. The barplots of CDR of QTN1 in each population under ten conditions: five levels of *F*_*ST*_ of QTN1 and two patterns of polymorphisms of QTN. Blue horizontal dashed lines indicate the CDR in the population A for each population. A: *japonica*, B: *indica*, C: temperate *japonica*, D: diverse, E: A+ B, F: A + C, G: A + D.

For almost all levels of *F*_*ST*_, the CDRs for QTN1 in the mixture populations E, F, and G were larger than in the corresponding non-mixture populations B, C, and D, regardless of the two QTN polymorphism patterns (Fig. 2). The CDRs for QTN1 in the mixture populations E, F, and G were always larger than in population A when all the simulation results were taken into account (Fig. 2). When *F*_*ST*_ was low, and all simulation results were taken into account (Fig. 2a), populations G and D, which were highly diverse populations, had a higher CDR than the other populations. When *F*_*ST*_ was in the lower-middle or middle category, and all simulation results were taken into account (Figs. 2b, c), population E had the highest CDR. The CDR of the highly diverse populations G and D significantly decreased as *F*_*ST*_ increased. This result suggested that the QTN1 effect could confound with the population structure at higher *F*_*ST*_ values, which meant that it was difficult to detect QTN1 in a highly diverse population. When the *F*_*ST*_ value was in the higher-middle or high level categories, and all simulation results were taken into account, (Figs. 2d, e), the CDR for QTN1 became quite low in all populations. In populations D, E, and G, QTN1 was hardly detected because of the strong confounding effect of the population structure. In the other populations, the expected heterozygosity (*H*_*e*_) for QTN1 was extremely small (In A and B, *H*_*e*_ was less than 0.1 in all 100 simulations). The small *H*_*e*_ may make the detection of QTN1 difficult.

We excluded the simulations in which there were no polymorphisms in the population so that the detection power of the GWAS when there were polymorphisms in an analyzed population could be evaluated (Figs. 2f-j). When *F*_*ST*_ was low, population F had the highest CDR and when *F*_*ST*_ was in the lower-middle or middle categories, population A had the highest CDR. However, there were only 14 and 9 cases in which QTN1 was polymorphic in population A. In general, the populations with low or moderate genetic diversities (A, C, and F) had higher CDRs than the populations with high genetic diversities (D, E, and G). When *F*_*ST*_ was in the higher-middle or high categories, the results were similar to when *F*_*ST*_ was in the lower-middle or middle categories.

The CDRs of QTN2 and QTN3 were much lower than that of QTN1 because smaller genetic variances were assigned to these QTLs than QTN1 (Table S2). As in the case of QTN1, for almost all levels of *F*_*ST*_, the CDRs of QTN2 and QTN3 were higher in the mixture populations (E, F, and G) than their corresponding non-mixture populations (B, C, and D). Furthermore, the CDRs for QTN2 and QTN3 in all the mixture populations were higher than for population A. The CDRs for QTN2 and QTN3 were also larger when the *F*_*ST*_ for QTN1 was higher.

Populations D and G had high AUC values in all cases (Table S2). Population F had a smaller AUC than populations D and G, even when the CDR was highest in population F.

### Comparisons of the −log_10_(*p*) values for the GWAS on each mixture population containing *japonica* (A)

We compared the −log_10_(*p*) values for each QTN between populations mixed with the *japonica* population (A) to see if QTN1 was polymorphic in A (Fig. 3). Comparing these values allowed us to examine whether the detection power of the GWAS improved when genetic resources with higher genetic diversities were added to target population A. There is no plot for the high *F*_*ST*_ values because no QTN1 was polymorphic in population A over 100 simulations when the *F*_*ST*_ of QTN1 was high.

**Fig. 3.**
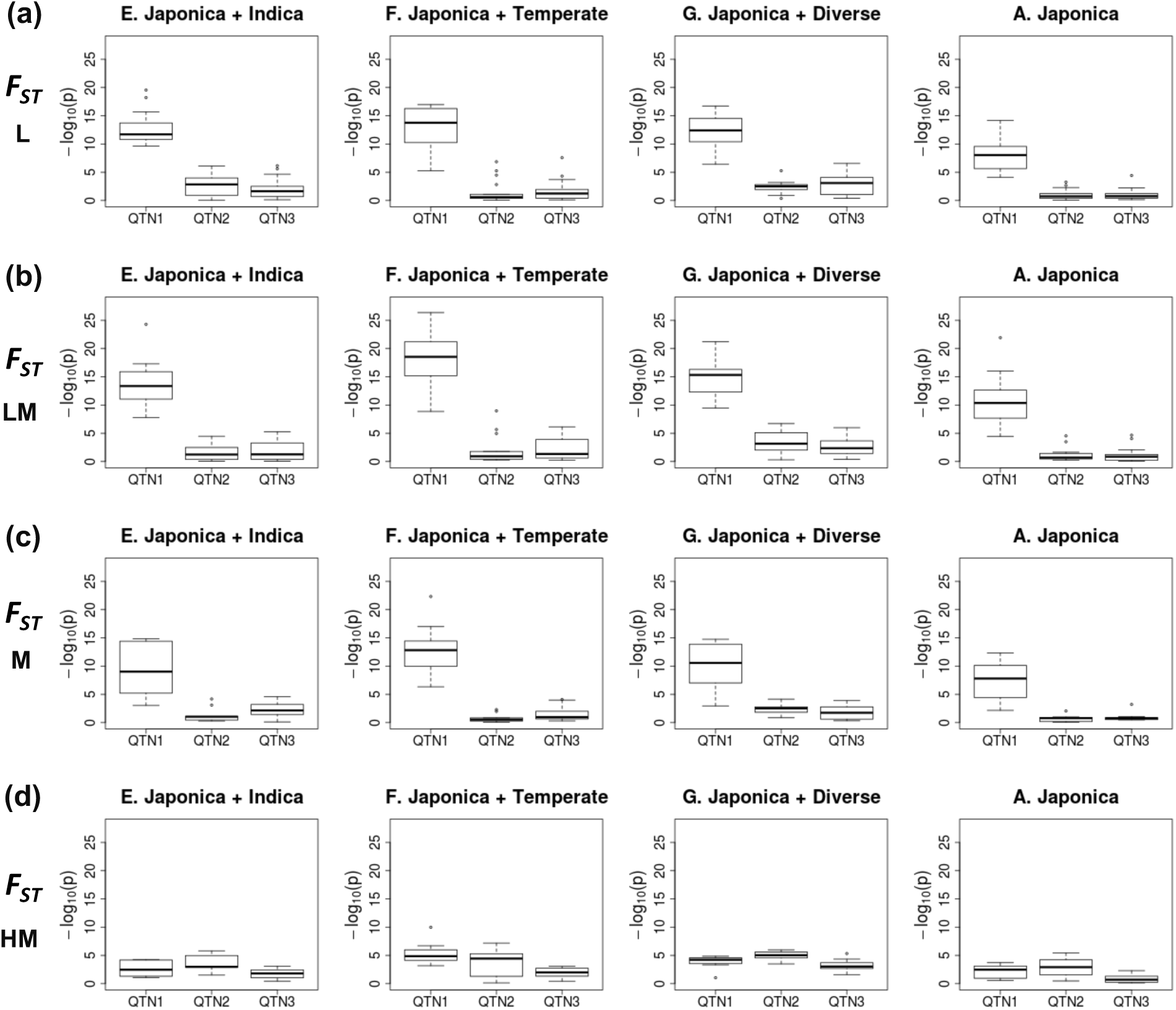
Boxplots of −log_10_(*p*) of each QTN when QTN1 was polymorphic in *japonica* (A). Boxplots of −log_10_(*p*) of each QTN for each mixture population and *japonica* (A) when QTN1 was polymorphic in A. These plots are shown divided into four categories according to the *F*_*ST*_ value for QTN1 (a: low, b: lower-middle, c: middle, d: higher-middle).

For all of the four *F*_*ST*_ levels, the detection power improved in all mixture populations compared to A (Fig. 3). Population F showed the highest detectability, and this tendency was conspicuous even when *F*_*ST*_ was in the middle or higher-middle categories (Figs. 3c, d, respectively). This is because the QTN1 effect is less likely to be confounded with the population structure in F than in the other mixture populations (E and G). Population G had the highest −log_10_(*p*) values for QTN2 and QTN3, although only slightly (Fig. 3).

### Factors affecting the detection power of QTNs in the mixture populations

We considered the factors related to the detection power of QTNs in the mixture populations by creating a figure that represented the relationship between *F*_*ST*_, the expected heterozygosity (*H*_*e*_), and the QTN1 detection power (Fig. 4 and Fig. S4 in Supplemental File 6).

**Fig. 4.**
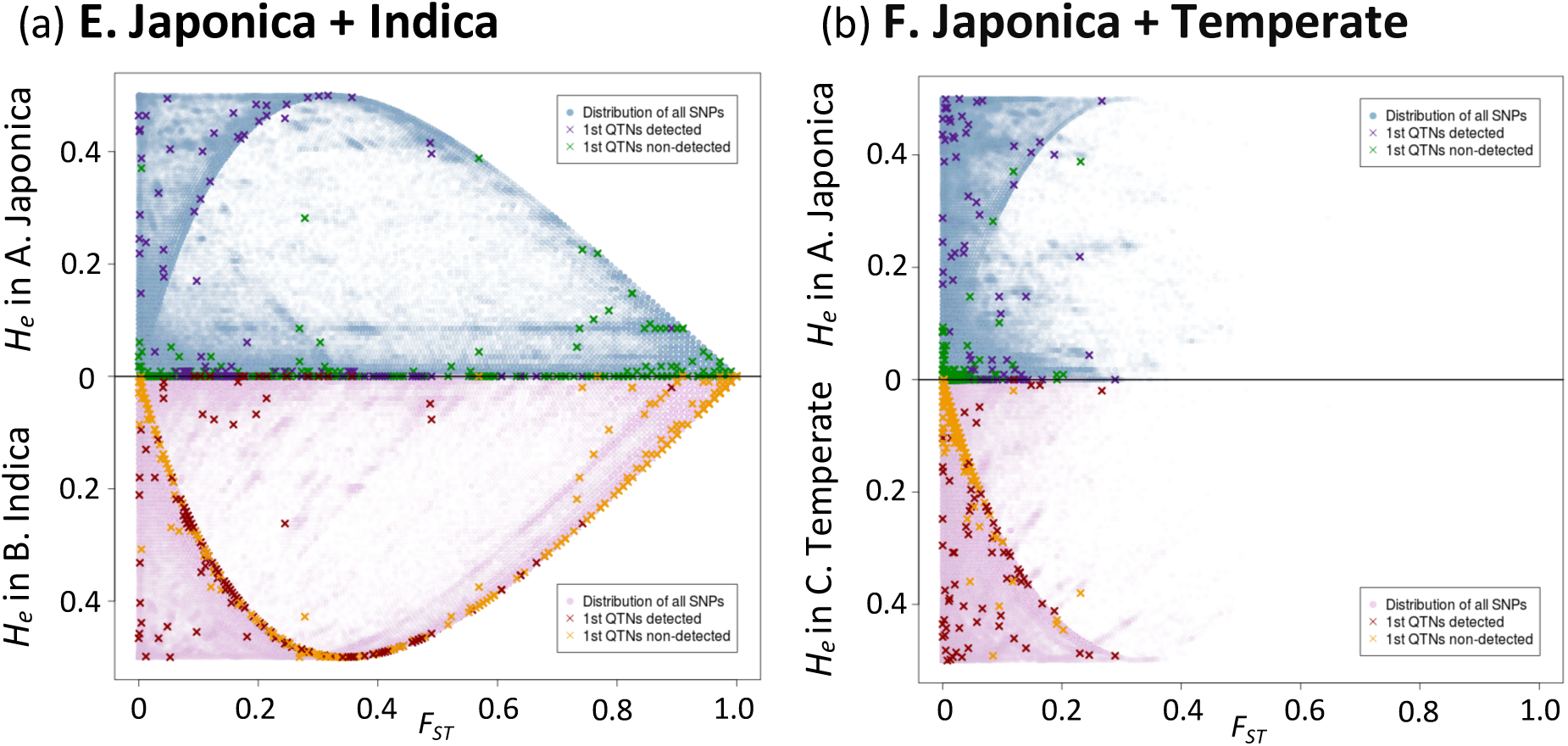
Relationship between *F*_*ST*_, *H*_*e*_, and the detection power of QTN1. The distribution of each marker is plotted thinly with between subpopulation *F*_*ST*_ on the horizontal axis and *H*_*e*_ of each subpopulation on the vertical axis. The dark X marks on the plot show the SNPs selected as QTN1s in this study. Red and purple marks were detected by GWAS, and green and yellow ones were not detected by GWAS.

Detection of the QTNs by the GWAS was generally difficult when the between-subpopulation *F*_*ST*_ value was high, or *H*_*e*_ was low (Fig. 4). There seemed to be a significant difference between plots F and E or G (Fig. 4a, Fig. S4, and Fig. 4b). However, in population F, because A and C are genetically close, the *F*_*ST*_ between the subpopulations was not high. Therefore, the relationships between *F*_*ST*_, *H*_*e*_, and the GWAS detection power applied to all mixture populations.

Some of the QTNs were detected by the GWAS when *F*_*ST*_ was in the medium category, and *H*_*e*_ in one of the subpopulations was close to 0 (Fig. 4a and Fig. S4). This suggested that even if the QTN was fixed in one subpopulation, the QTN may still be detected by the GWAS if another subpopulation was added to the analysis.

### Comparisons among the precision, recall, and *F*-measure values for each population

The three summary statistics (the mean of precision, the mean of recall, and the mean of the *F*-measure) were also calculated under ten conditions (Fig. S5 in Supplemental File 7). The precision of the mixture populations was better than the precision value for population A for almost all *F*_*ST*_ categories when all simulation results were taken into account (Fig. S5a). However, it is not necessarily true that the precision of the mixture populations outperformed that of their original genetic resources (compare E with B, F with C, and G with D). The recall values of the mixture populations were larger than for their original genetic resources under all conditions. Finally, a comparison of the *F*-measure for each population showed that there seemed to be no tendency associated with *F*_*ST*_. Therefore, it was difficult to conclude which population was suitable for a GWAS when the *F*-measure is used. These results indicated that using mixture populations for a GWAS led to the detection of more SNPs, including QTNs.

### Relationship between the CDR results and genetic diversity

The relationship between the CDR results for QTN1 and the degree of genetic diversity was evaluated under the two QTN polymorphism patterns, i.e., whether or not QTN was polymorphic in the population (Fig. S6 in Supplemental File 8). The CDRs for the mixture populations were usually larger than for the non-mixture populations if their diversity levels were close (Fig. S6a, b). A comparison of the results for the different *F*_*ST*_ categories showed that when *F*_*ST*_ was low, the populations with the highest diversities, such as D or G, had the highest CDRs, and when *F*_*ST*_ was in the lower-middle or middle categories, the populations with the second-highest diversities, such as B or E, had the highest CDRs. Finally, when *F*_*ST*_ was in the higher-middle or high categories, the populations with relatively low diversities, such as C or F, had the highest CDRs (Fig. S6a). However, when the simulations in which there were no polymorphisms in the population were excluded, the populations with relatively low diversities, such as A, C, or F, had the highest CDRs in almost all the *F*_*ST*_ categories (Fig. S6b).

## DISCUSSION

### Relationship between *F*_*ST*_ and QTN detection

One of the main results of this study was that the detection of QTNs was difficult in populations with high genetic diversities, such as D, E, and G, when the *F*_*ST*_ for QTN1 between *japonica* (A) and *indica* (B) was high. This was because the QTN effect confounds with the effect of population structure in these populations. We also examined the reasons why the CDRs for QTN2 and QTN3 were high when the QTN1 *F*_*ST*_ value was high.

In this study, phenotypic values were simulated using the following expression:

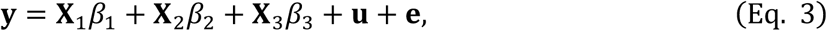

where **u** is the polygenetic effect, and is the term that reflects differences between accessions and thus differences between subpopulations. Therefore, if the degree of QTN1 genetic differentiation between *japonica* (A) and *indica* (B) is high, it can be assumed that there is a high correlation between **X**_1_*β*_1_ and **u**. In this study, we generated phenotypic values using a certain variance ratio under the assumption that each term is independent. Therefore, if there is a correlation between **X**_1_*β*_1_ and **u**, and the variance between these two terms is considered as one unit, it can be assumed that the variance is smaller than the total value of the two variances under the assumption of independence. Therefore, the variance of these two terms (**X**_1_*β*_1_ + **u**) in the total phenotypic variance becomes smaller, whereas the variances caused by the terms **X**_2_*β*_2_ and **X**_3_*β*_3_ become greater than those when it is assumed that each term is independent.

The GWAS model used in this study was

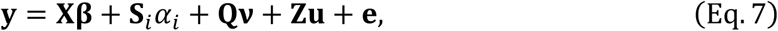

where **Q*v*** is the term used to correct the effect of population structure, and Zu shows the polygenetic effect. In this GWAS model, **S**_*i*_*α*_*i*_ and **Q*v*** or **Zu** have some correlation when **S**_*i*_ = **X**_1_. This correlation results in the underestimation of *α*_*i*_ by the terms originally used to correct the effects of population structure or family relatedness, such as **Q*v*** and **Zu**. Therefore, QTN detection is quite difficult when a GWAS is performed on mixture populations. For QTN2 and QTN3, where **S**_*i*_ = **X**_2_ or **S**_*i*_ = **X**_3_, there is generally no correlation between **S**_*i*_*α*_*i*_ and **Q*v*** or **Zu**. Therefore, the detection of these QTNs is not related to these terms. Furthermore, the variances represented by the terms **X**_2_*β*_2_ and **X**_3_*β*_3_ are considered to be higher when the QTN1 genetic differentiation is not high. Therefore, the CDRs of QTN2 and QTN3 were high when the *F*_*ST*_ for QTN1 was high (Fig. 2 and Table S1). It has been suggested by Atwell et al. (2010) that a bias may occur in the GWAS results when the QTN correlates with population structure or family relatedness.

### Relationship between *H*_*e*_ and the QTN detection

The detection of QTNs by a GWAS was difficult when the expected heterozygosity (*H*_*e*_) in the population was low. When *H*_*e*_ in the population was low, the MAF was low, and alleles and mutations with low allele frequencies are known as “rare alleles” or “rare variants”. In such cases, the QTN effect when the *H*_*e*_ values are low may be concealed by the QTN effect when *H*_*e*_ is not low or by the environmental effect because there are few accessions with one allele. For this reason, it is generally challenging to detect QTNs in such cases, but a method to deal with this problem has been developed (Wu et al., 2011).

One example of a “rare variant” that is common in plants is the haplotype condition. It has been reported that haplotypes are difficult to detect using a GWAS (Stram, 2014). This is because haplotypes are often “rare variants” and their *H*_*e*_ values in the population are low. Another problem caused by “rare variants” is that the non-causal SNP whose LD is strong with the “rare variant” may have a higher −log_10_(*p*) value than the “rare variant”. This occurrence, known as “synthetic association”, often happens when the minor allele frequency of the SNP is higher than that of the “rare variant” (Dickson et al., 2010). These “synthetic associations” were often detected in this simulation study.

### Summary and further discussion on each result

Generally, the CDRs of the QTNs showed that the populations suitable for a GWAS were different depending on whether all the QTNs were to be detected or only the polymorphic QTNs in the target population. Specifically, if all QTNs are to be detected when the degree of genetic differentiation between QTNs is low, then it is optimal to use a population with high genetic diversity that has as many polymorphisms as possible. However, as the degree of genetic differentiation becomes more extensive, a population with high genetic diversity is not suitable for a GWAS because the QTN effect is more likely to confound with the population structure. In contrast, a population with moderate genetic diversity, such as population F, was suitable for a GWAS, regardless of the degree of genetic differentiation. This was partly because the QTN1 effect was less likely to confound with the population structure in F than in E or G, even when *F*_*ST*_ was high. However, in either case, when the degree of genetic differentiation is extensive, it is difficult to detect the QTNs in any population. Therefore a GWAS analysis is not suitable, which means that another approach, such as biparental QTL mapping, must be used to identify genes (Lander and Botstein, 1989).

Population F had a smaller AUC than populations D and G, even when the CDR for population F was the highest. From its definition, AUC is more dependent on how low −log_10_(*p*) of the QTN with the lowest −log_10_(*p*) value is than on how high the −log_10_(*p*) of the QTN with the highest −log_10_(*p*) value is. Furthermore, in this study, the number of markers for the GWAS differed (Table 1). When −log_10_(*p*) values for the QTNs were similar among the different populations, the larger number of markers meant that the true negative rate increased, and the false positive rate decreased in a population, which resulted in an increase in the AUC of a population with a larger number of markers, e.g. D and G.

A comparison of the mixture populations and *japonica* (A) using −log_10_(*p*) showed that when the QTNs are polymorphic in a target population with low genetic diversity, genetic resources with higher genetic diversities should be added to the target population. However, in order to avoid cases where the degree of genetic differentiation among the QTNs is extensive between the target population and genetic resources, it is desirable to use populations that are genetically close to the target population.

Finally, the results suggested that the *F*_*ST*_ differences between the subpopulations and the expected heterozygosity (*H*_*e*_) of each subpopulation greatly influenced QTN detection by the GWAS in the mixture populations (Fig. 4 and Fig. S4). This result was in agreement with the above finding that QTN detection using a GWAS was generally difficult when *F*_*ST*_ was high, or *H*_*e*_ were low. However, these situations frequently happened when the *F*_*ST*_ between the subpopulations was moderate. Therefore, even if a QTN is fixed in one subpopulation, it may be possible to detect the QTN by adding another population to the analysis because when the *H*_*e*_ of the QTN is low in one population and *F*_*ST*_ is moderate, it can be assumed that *H*_*e*_ is relatively high in the other population. Therefore, the *H*_*e*_ of the mixture population as a whole becomes larger and the detection of a QTN is possible unless the confounding of the effect of that QTN with the population structure is extensive. Although this situation is not difficult to interpret, it is extremely important that SNPs with high *F*_*ST*_ and low *H*_*e*_ values must exist in large numbers among populations. After taking this fact into account, a GWAS with a mixture population can be useful. Therefore, creating the proposed diagram shown in Fig. 4 and Fig. S4, will lead to a quantitative understanding of what kind of SNPs can be detected by a GWAS in mixture populations of interest.

### Relationships with using whole-genome sequences

One of the major factors related to the QTN detection power was the fixation index *F*_*ST*_ differences among subpopulations. When the *F*_*ST*_ difference between the *japonica* (A) and *indica* (B) subpopulations was low, the CDR of the mixture populations was high. One example of such markers is that mutations may have occurred at the same position in both populations after they differentiated. Since such variants are relatively new variants, the LD relationship between these variants and surrounding markers will be weak. Therefore, these variants cannot be detected using marker genotype data with a small number of markers, such as an SNP array. However, the use of whole-genome sequences will increase the marker density, which improves the possibility of detecting such variants with a GWAS. In summary, using whole-genome sequences improves the possibility of detecting QTNs with low *F*_*ST*_ values and the use of mixture populations should further improve the QTN detection power. In this study, there were cases where SNPs in a low LD region were selected as QTNs when *F*_*ST*_ was low.

## CONCLUSION

In this study, we examined a way of selecting a population that was suitable for a GWAS by conducting simulations using populations with various genetic backgrounds. We evaluated the results of the simulations by dividing them into ten patterns according to two criteria: the degree of genetic differentiation (*F*_*ST*_) between two main subpopulations and QTN polymorphism in a target population. When the QTNs are polymorphic in a target population, increasing the population size by adding available genotypes to the target population improves the detection power. We suggest that a population genetically similar to a target population is desirable. After investigating *F*_*ST*_ and expected heterozygosity *H*_*e*_ as factors that may substantially influence the detection power of a GWAS, the results showed that SNPs with high *F*_*ST*_ and low *H*_*e*_ values were less likely to be detected by a GWAS that used mixture populations. These results indicated that the detection power of a GWAS was improved by using mixture populations with different genetic backgrounds. Furthermore, the use of publicly available whole-genome sequences meant it was possible to increase the population size and to use polymorphic markers that were present in high numbers, which should also improve the detection power of the GWAS.

## ACKNOWLEDGMENTS

This study was supported by Grant-in-Aid for Scientific Research(A) (25252002), Grant-in-Aid for Scientific Research(B) (Grant number 15H04436), JST, PRESTO (Grant number JPMJPR15O6), the Cross-ministerial Strategic Innovation Promotion Program (SIP), the “Technologies for creating next-generation agriculture, forestry and fisheries” (funding agency: Bio-oriented Technology Research Advancement Institution, NARO), and JST CREST (Grant Number JPMJCR16O).

We are also grateful for the advice given by Dr. Ryokei Tanaka.

## SUPPLEMENTAL MATERIAL

Supplemental File 1: **Table S1.** Information about the 299 rice accessions used in this study.

Supplemental File 2: **Fig. S1.** Histogram showing the *F*_*ST*_ differences between *japonica* (A) and *indica*

(B).

Supplemental File 3: **Fig. S2.** Example of a ROC curve and the AUC.

Supplemental File 4: **Fig. S3.** Principal components analysis results for chromosome 1 and all the chromosomes.

Supplemental File 5: **Table S2.** Correct detection rate rates for all QTNs and the AUC in each population.

Supplemental File 6: **Fig. S4.** Relationship between *F*_*ST*_, *H*_*e*_, and the QTN1 detection power for the population G.

Supplemental File 7: **Fig. S5.** Bar plots of the precision, the recall and the *F*-measure results.

Supplemental File 8: **Fig. S6.** Relationship between the diversity level and the CDR of QTN1.

Supplemental File 9: **Supplementary Note.** Additional information about the materials used in this study.

## OPTIONAL SECTIONS

### Availability of data and material

Whole genome sequencing data are available of 112 accessions of *Oryza sativa* subsp. *japonica* in the DDBJ Sequence Read Archive (SRA106223, ERA358140, DRA000158, DRA000307, DRA000897, DRA000927, DRA007273, DRA007256, and DRA008071). Whole genome sequencing data for all the other accessions are available in the “Rice SNP-Seek Database”.

### Competing interests

The authors declare that they have no competing interests.

### Author’s contributions

KH, HKK, and HI conceived and designed the study. KH and HI performed the mathematical and statistical analysis. KH, HKK, MY, EK, SY and HN contributed to marker genotyping. KH, HKK, and HI wrote the manuscript in consultation with MY, EK, SY, and HN. All authors read and approved the final manuscript.

