## Supplementary figures and images for "Choosing the optimal population for a genome-wide association study: a simulation using whole-genome sequences from rice"

### Supplemental Figure 1

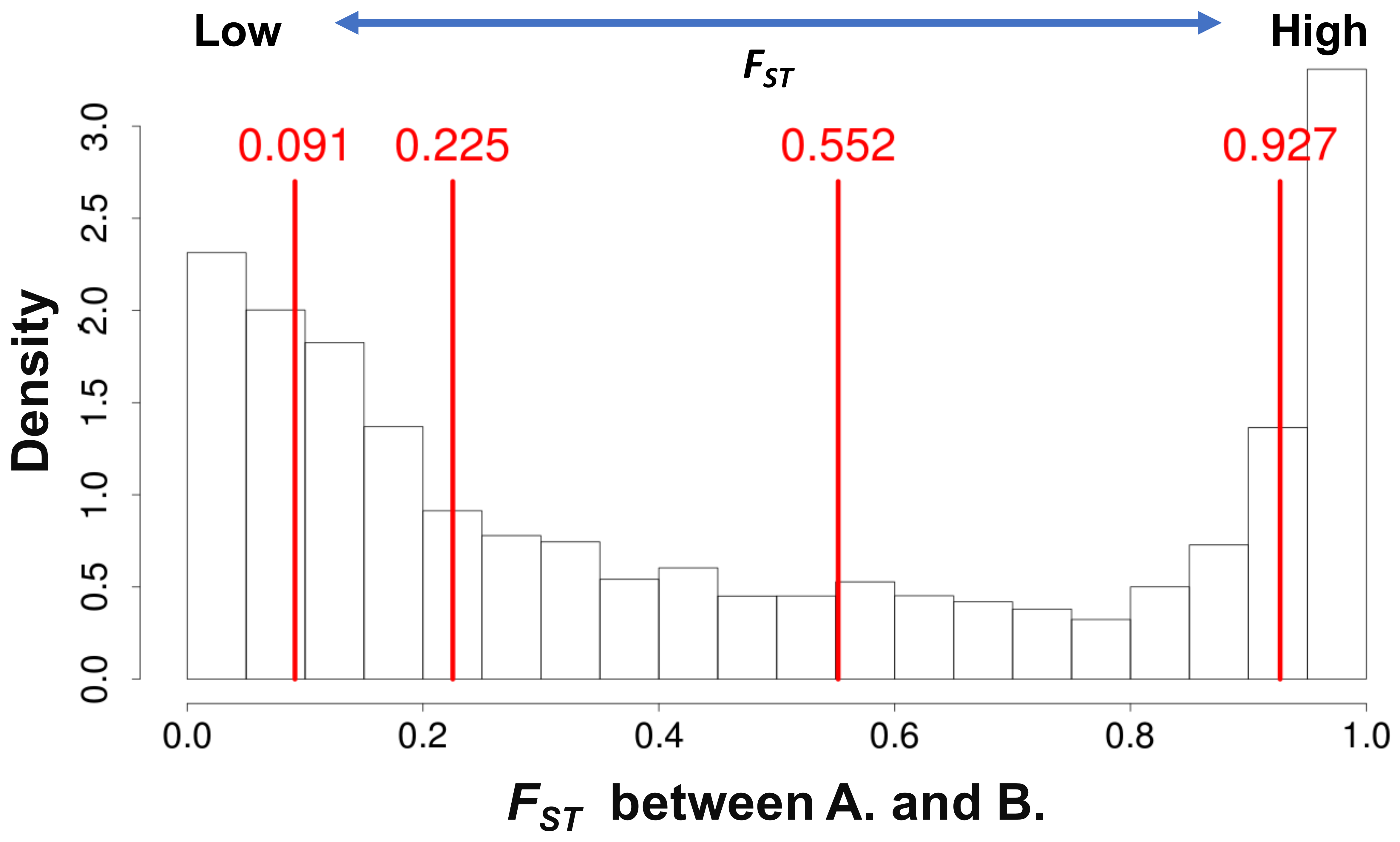

### Supplemental Figure 2

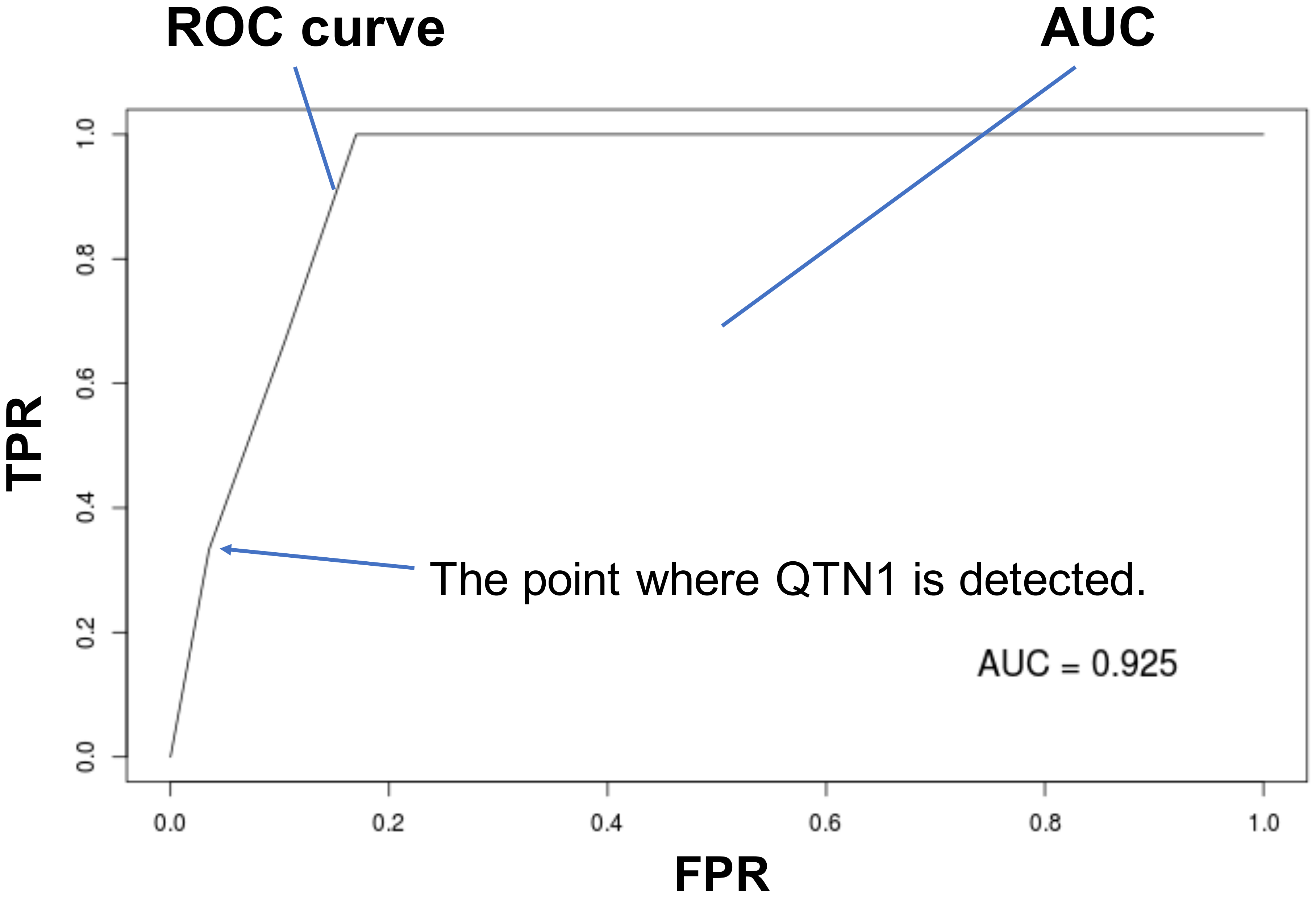

### Supplemental Figure 3

##### The result of PCA for chromosome 1

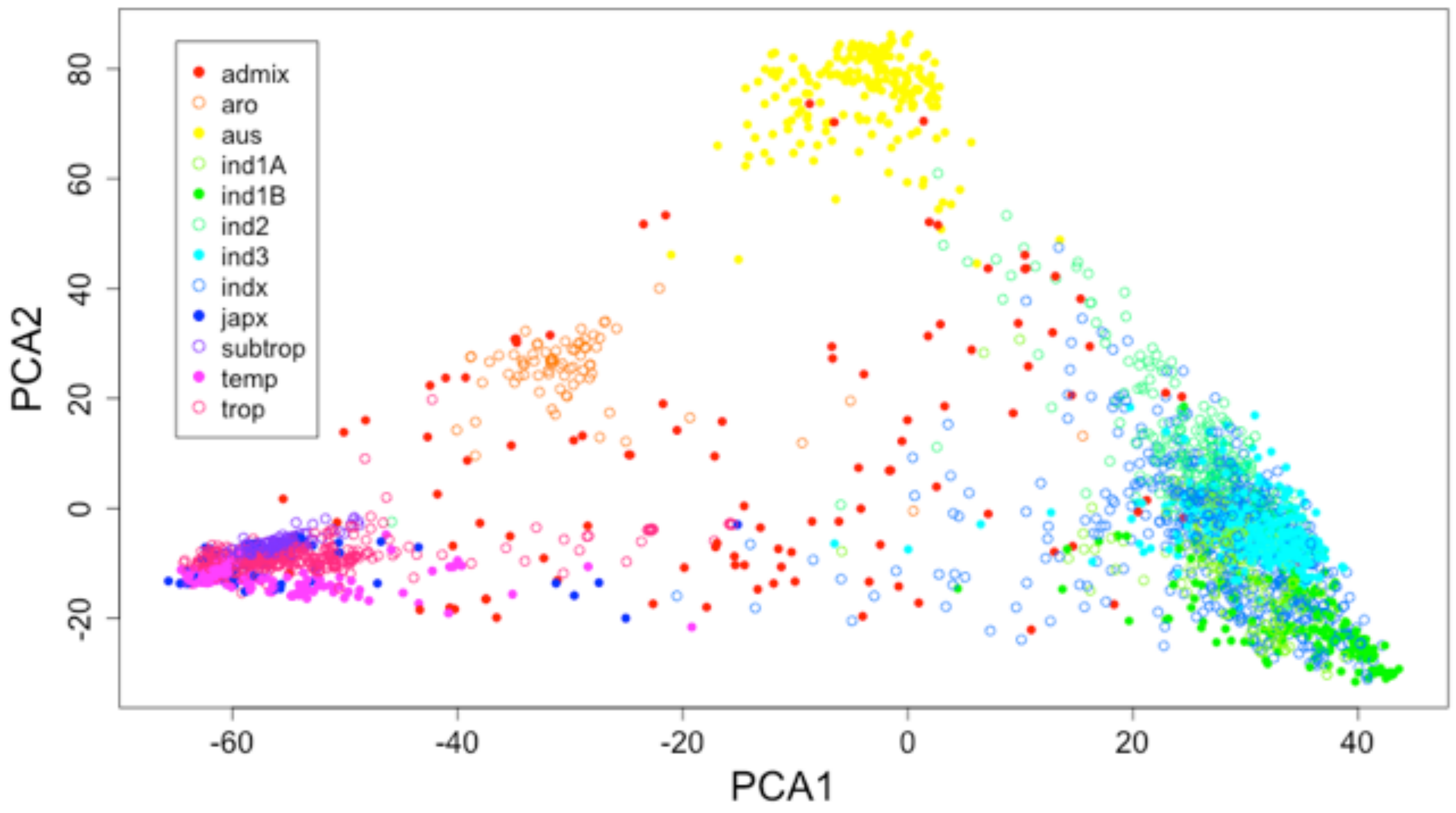


##### The result of PCA for all chromosome

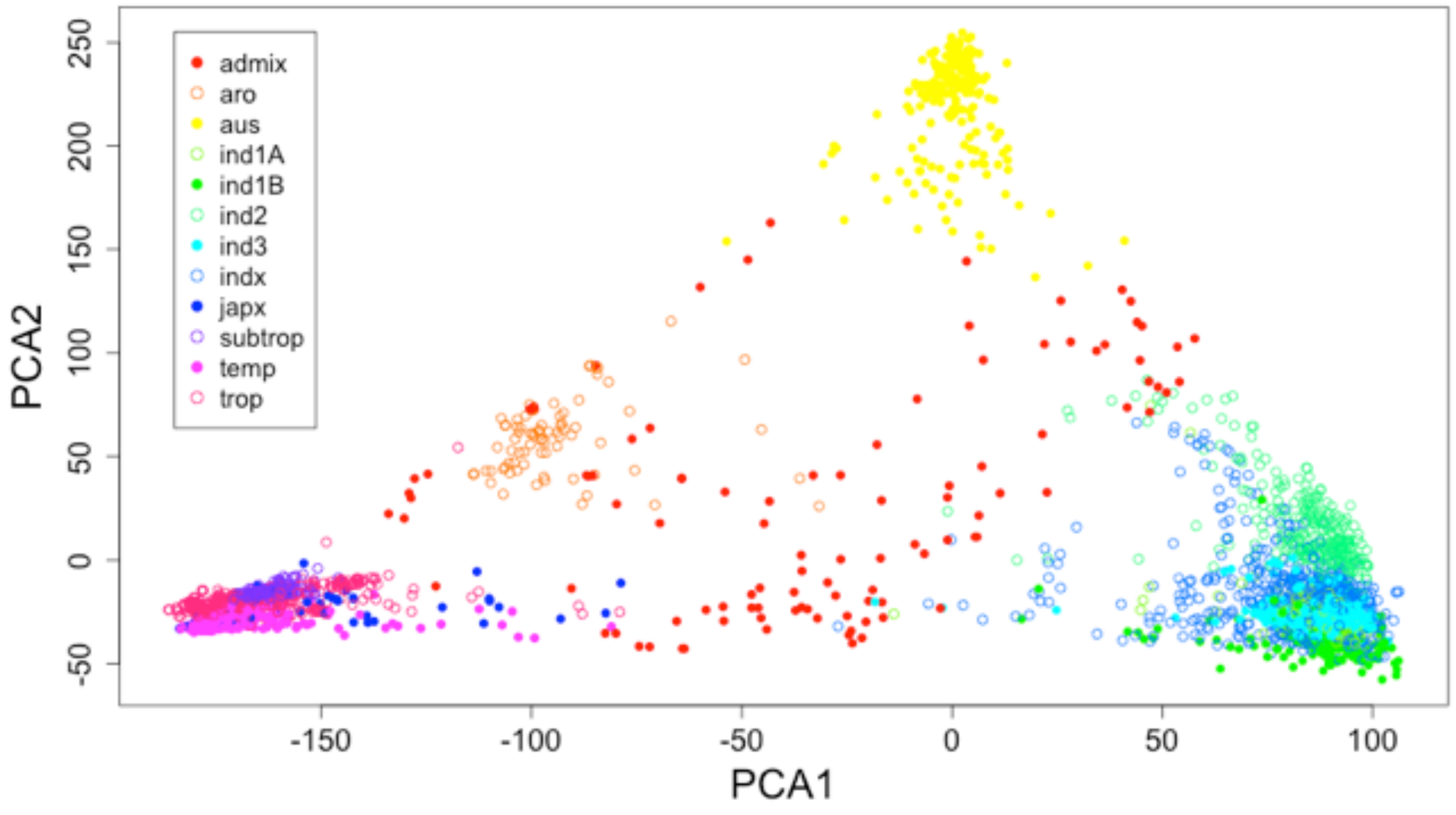

### Supplemental Figure 4

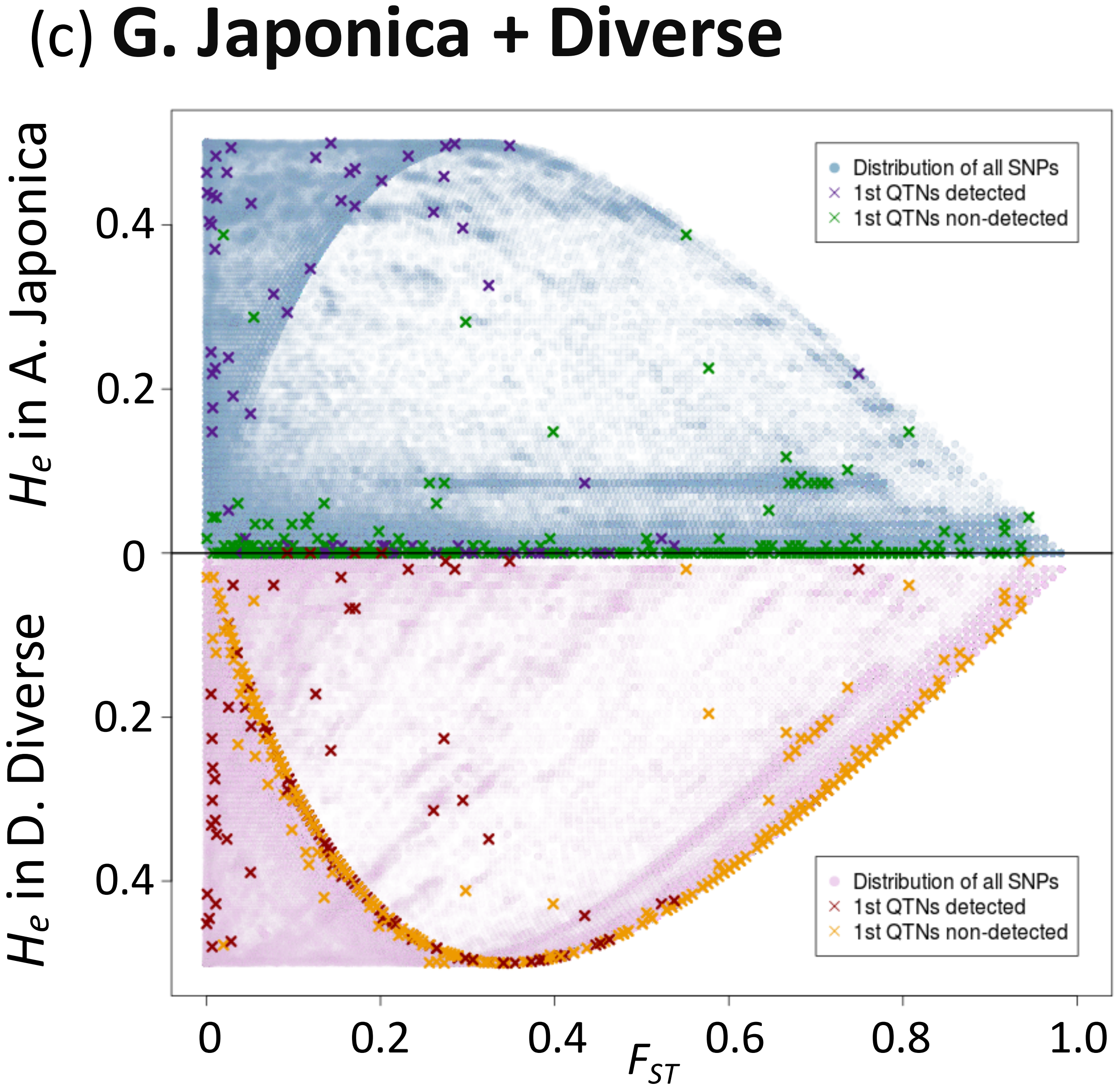

### Supplemental Figure 5

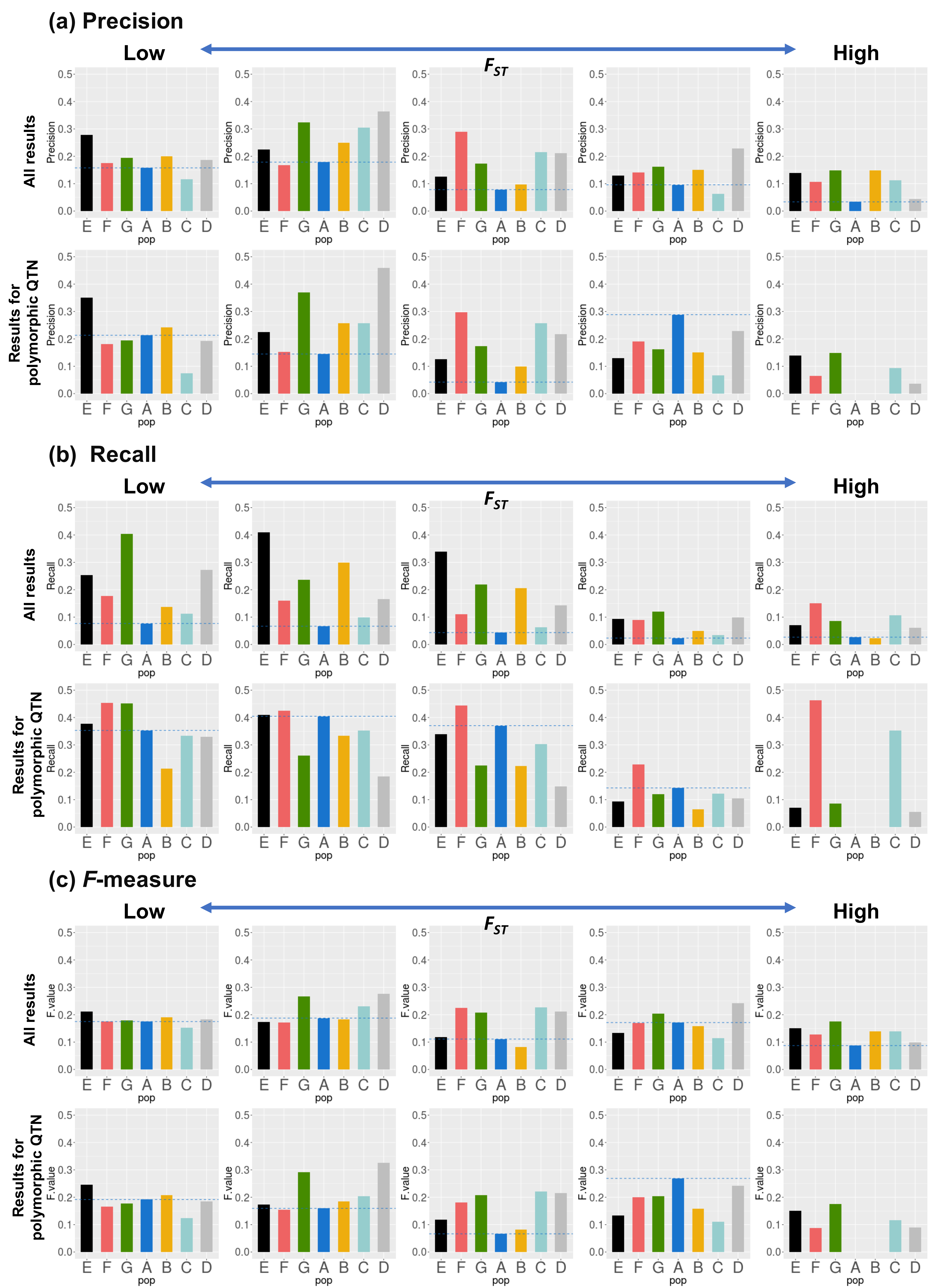

### Supplemental Figure 6

#### (a) All results

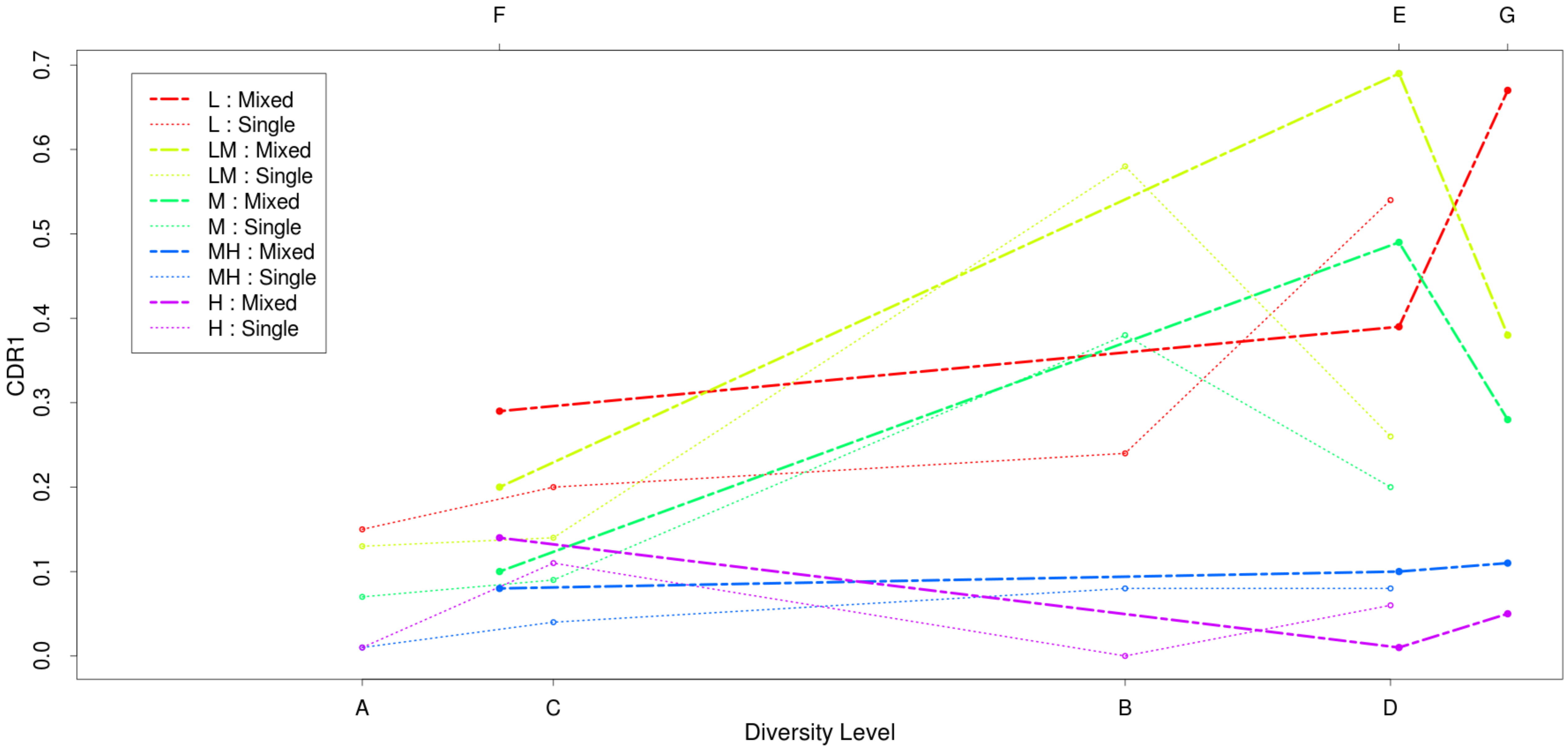


#### (b) Results for polymorphic QTN

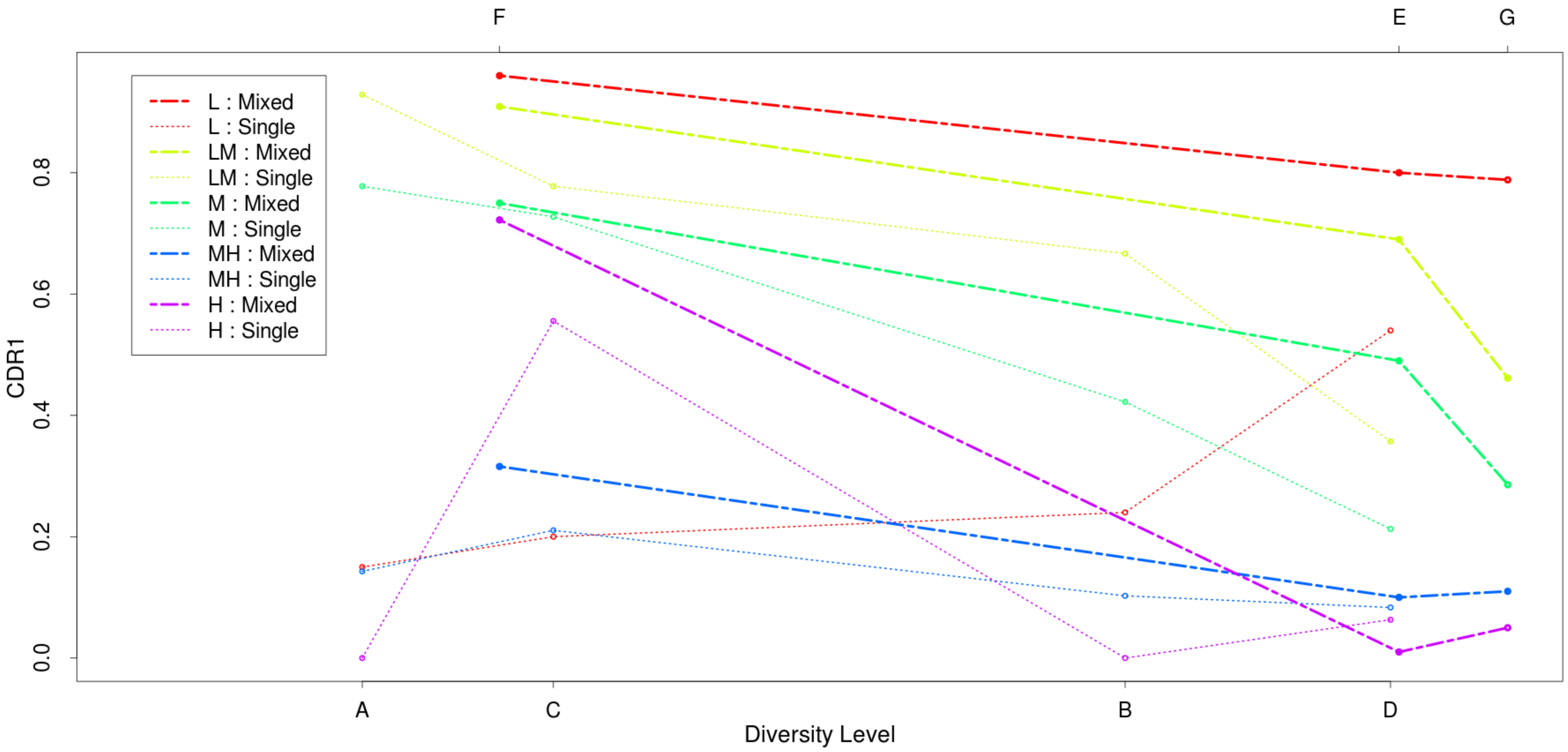
