## Supplemental Note 1 for "Choosing the optimal population for a genome-wide association study: a simulation using whole-genome sequences from rice"

**Additional information about the materials**

On the “Rice SNP-Seek Database” website, marker genotype data with 3,024 accessions of rice are available. These 3,024 accessions are categorized into 12 subpopulations (“admix”, “aro”, “aus”, “ind1a”, “ind1b”, “ind2”, “ind3”, “indx”, “japx”, “subtrop”, “temp”, and “trop”). In this study, 100 accessions of B were selected from 209 accessions whose subpopulation is “ind1a”, and 100 accessions of C were selected from 288 accessions whose subpopulation is “temp”, and 100 accessions of D were selected from 3,024 accessions regardless of subpopulations. These populations were selected in the same manner, using the “k-medoids” method. Original accessions from the “Rice SNP-Seek Database” website (209, 288 and 3,024 respectively) were classified into 100 groups by k-medoids, and then 100 medoids, which represented each class, were selected as accessions representing the original populations of the “Rice SNP-Seek Database” website. These selection were performed based on the marker genotype data for 3,024 accessions consisting of core SNPs (defined by Rice SNP-Seek Database as “404K CoreSNP Dataset”) of all 12 chromosomes, using the “pam” function of the R package “cluster” version 2.0.9 (Maechler et al., 2018[1]).

In this study, the marker genotype data of chromosome 1 of the whole-genome sequences were used for GWAS. We emphasize that there is almost no issue that not the marker genotype data consisting of all chromosomes, but that of only chromosome 1 were used for GWAS. We performed the principal component analysis for the marker genotype data consisting of all chromosomes and that of only chromosome 1, then compared these results (Figure S3 in Additional file 4). The correlations of the first and second principal components between them were 0.988 and 0.957, respectively. Therefore, we concluded that the marker genotype data of chromosome 1 represented the nature of the marker genotype data with all chromosomes.
